# The Rbfox1/LASR complex controls alternative pre-mRNA splicing by recognition of multi-part RNA regulatory modules

**DOI:** 10.1101/2024.07.12.603345

**Authors:** Parham Peyda, Chia-Ho Lin, Kelechi Onwuzurike, Douglas L. Black

## Abstract

The Rbfox proteins regulate alternative pre-mRNA splicing by binding to the RNA element GCAUG. In the nucleus, most of Rbfox is bound to LASR, a complex of RNA-binding proteins that recognize additional RNA motifs. However, it remains unclear how the different subunits of the Rbfox/LASR complex act together to bind RNA and regulate splicing. We used a nuclease-protection assay to map the transcriptome-wide footprints of Rbfox1/LASR on nascent cellular RNA. In addition to GCAUG, Rbfox1/LASR binds RNA containing motifs for LASR subunits hnRNPs M, H/F, C, and Matrin3. These elements are often arranged in tandem, forming multi-part modules of RNA motifs. To distinguish contact sites of Rbfox1 from the LASR subunits, we analyzed a mutant Rbfox1(F125A) that has lost RNA binding but remains associated with LASR. Rbfox1(F125A)/LASR complexes no longer interact with GCAUG but retain binding to RNA elements for LASR. Splicing analyses reveal that in addition to activating exons through adjacent GCAUG elements, Rbfox can also stimulate exons near binding sites for LASR subunits. Mini-gene experiments demonstrate that these diverse elements produce a combined regulatory effect on a target exon. These findings illuminate how a complex of RNA-binding proteins can decode combinatorial splicing regulatory signals by recognizing groups of tandem RNA elements.

## Introduction

Eukaryotes produce multiple mRNAs from the same pre-mRNA by changing the choice of splice sites that define an exon. This process of alternative splicing is regulated by trans-acting RNA-binding proteins (RBPs) that bind to cis-regulatory elements on pre-mRNAs (Black 2003). RBPs can either enhance or repress exon inclusion. Their effects depend on their binding position relative to the target exon and on the actions of other factors binding nearby. Alternative exons usually carry binding elements for multiple RBPs that potentially interact with each other, giving rise to a complex combinatorial code that is difficult to unravel (Smith and Valcárcel 2000; Ule and Blencowe 2019).

One family of regulators controlling important splicing programs in the nervous system and over development is the Rbfox proteins (Conboy 2017). These RBPs, conserved from *C. elegans* to humans, are unusual for having a single RNA Recognition Motif (RRM) that is highly specific for the element GCAUG (Auweter et al., 2006; Begg et al., 2020; Jin et al., 2003; Lambert et al., 2014; Ponthier et al., 2006; Underwood et al., 2005; Ye et al., 2023). This sequence specificity is due to an unusual interaction with the RRM domain that includes many protein contacts along the short motif (Auweter et al. 2006). Mammals have three Rbfox genes: *RBFOX1* is abundant in the brain, heart, and muscle, *RBFOX2* exhibits broad expression across tissues during development, and *RBFOX3* appears exclusive to the brain (Conboy 2017). Each of these genes produces multiple spliced products, including cytoplasmic isoforms that regulate translation and other processes, and nuclear isoforms that regulate splicing (Carreira-Rosario et al. 2016; Damianov and Black 2010; Lee et al. 2016; Vuong et al. 2018). Besides its conserved RNA binding domain, each Rbfox also contains a low complexity, tyrosine-rich C-terminal domain (CTD). This CTD can homo-oligomerize and also binds to the large assembly of splicing regulators (LASR), a heteromeric complex of RBPs consisting of hnRNP M, hnRNP H/F, hnRNP C, DDX5, NF-110, hnRNP UL2, NF-45, Matrin3, and in some preparations MeCP2 (Damianov et al. 2016; Jiang et al. 2021; Ying et al. 2017). The repetitive tyrosines within the CTD are required for assembly of Rbfox/LASR into higher-order complexes and its ability to activate splicing (Ying et al. 2017).

Rbfox proteins play critical roles in the development and function of multiple organs. In the central nervous system, mutation or aberrant expression of these proteins can lead to electrophysiological abnormalities, seizures, and defects in cerebellar development in mice, and to epileptic and/or autism spectrum disorders in human patients (Gehman et al. 2011, 2012; Jacko et al. 2018; Lal et al. 2013a, 2013b; Vuong et al. 2018). In the heart, dysregulation of Rbfox2 can lead to hypoplastic left heart syndrome, congenital heart disease, and conduction defects in Myotonic Dystrophy 1 (Homsy et al., 2015; Huang et al., 2022; Misra et al., 2020; Verma et al., 2016). Rbfox2 is also involved in cholesterol homeostasis in the liver and controlling insulin secretion from the pancreas (Moss et al. 2023; Paterson et al. 2022). Moreover, Rbfox2 contributes to epithelial-mesenchymal transition and is important for pancreatic cancer metastasis and breast cancer development (Jbara et al. 2023; Li et al. 2018; Maurin et al. 2023; Shapiro et al. 2011; Venables et al. 2013).

*In vivo* binding sites for RBPs can be identified using crosslinking and immunoprecipitation (CLIP) (Hafner et al., 2021). Multiple studies have correlated Rbfox crosslinked sites with exons whose splicing is also regulated by the protein (Begg et al. 2020; Damianov et al. 2016; Jangi et al. 2014; Lovci et al. 2013; Weyn-Vanhentenryck et al. 2014; Yeo et al. 2009). These analyses confirmed that all three Rbfox proteins bind GCAUG, with a preference for UGCAUG, while close variants of this motif also crosslinked. Binding to GCAUG downstream of an exon was found to usually enhance its inclusion while upstream binding was associated with exon repression. However, GCAUG is a common pentamer in genomic RNA, and not all GCAUGs in the transcriptome crosslink to Rbfox. Conversely, some Rbfox crosslinked sites do not contain GCAUG or its related secondary motifs (Begg et al. 2020; Damianov et al. 2016; Jangi et al. 2014; Weyn-Vanhentenryck et al. 2014; Yeo et al. 2009). These sites might contain undefined motifs with an affinity for Rbfox or could arise from Rbfox crosslinking by virtue of its proximity to a sequence without a specific interaction.

Rbfox’s association with LASR helps explain its selectivity for particular GCAUG elements (Damianov et al. 2016; Ying et al. 2017). GCAUGs that crosslink to Rbfox were found to be enriched for adjacent elements predicted to bind the LASR subunit hnRNP M, suggesting cooperative binding. Furthermore, many Rbfox crosslinked sites lacking GCAUG contain predicted binding motifs for the LASR subunits hnRNP M and hnRNP C. An analysis of CLIP for Rbfox, hnRNP M, hnRNP C, and SRSF1 indicated that Rbfox may regulate targets through direct binding, cooperative binding with a partner, or indirect binding via a partner (Zhou et al. 2021). However, it remains unclear if these effects result from Rbfox co-binding with LASR and what segments of RNA get bound by the entire Rbfox/LASR complex.

In this study, we map the transcriptome-wide RNA footprints of Rbfox1/LASR in HEK293 cells using a subcellular fractionation and nuclease-protection assay. These footprints provide information on the extent of RNA contact by the protein complex at different regulatory sites. We find that these regions consist of multi-part RNA elements that can bind Rbfox and the LASR subunits hnRNP M, hnRNP H/F, hnRNP C, and Matrin3. Comparing binding of LASR containing wildtype Rbfox1 with that containing the RNA-binding mutant F125A allowed us to distinguish the binding sites of Rbfox1 from those of LASR subunits. By comparing splicing changes induced by wildtype and F125A Rbfox1 in cells, we identify exons requiring Rbfox binding to GCAUG, as well as exons regulated by Rbfox secondary motifs and LASR subunit motifs. Mini-gene experiments indicate that both GCAUG and other elements combined in tandem exert positive effects on exon inclusion.

## Results

### Mapping Rbfox1/LASR binding sites on chromatin-associated RNA by nuclease-protection

We recently reported IP-seq as a method to map the sites of U2 snRNP binding across the transcriptome (Damianov et al. 2024). This approach involves isolation of ribonucleoprotein (RNP) complexes from the chromatin fraction of cells after extraction with nuclease. We found that the isolated U2 snRNP in this fraction was bound to protected RNA fragments corresponding to branchpoints basepaired to the U2 snRNA. This method can potentially map the contact sites of other factors bound to nascent RNA, provided their binding affinity is sufficient to withstand the nuclease degradation. Using the same chromatin extraction, we previously isolated FLAG-tagged Rbfox proteins from HEK293 cells and examined their interacting protein partners. We found that nuclear Rbfox proteins were almost entirely bound by the large assembly of splicing regulators, LASR, a complex of other RBPs (Damianov et al., 2016). Given the results with U2 snRNP, we examined whether there were nuclease-protected RNAs within the Rbfox/LASR preparations.

We engineered a cell line to express FLAG-tagged Rbfox1. Rbfox1 and Rbfox3 are not expressed in HEK293 cells, and we previously created a HEK293 cell line with the endogenous Rbfox2 knocked out (Damianov et al. 2016). We used this Rbfox deficient line as a recipient for a doxycycline inducible FLAG-Rbfox1 construct whose expression could be titrated to levels found in the brain. From these cells, we isolated nuclei, lysed them with Triton X-100, and centrifuged the lysate to obtain a soluble nucleoplasmic supernatant and a pellet containing chromatin and other high molecular weight material (Fig. 1A). The chromatin pellet was treated with Benzonase nuclease, degrading both RNA and DNA, to solubilize material within the pellet. Anti-FLAG immunoprecipitates were isolated from both the nucleoplasm and the chromatin extract, eluted with FLAG peptide, and analyzed by SDS-PAGE stained for total protein (Fig. 1B). As seen previously, Rbfox1 was more abundant in the chromatin extract than the soluble nucleoplasm. This FLAG-Rbfox1 copurified with additional bands of roughly equal intensity corresponding to the LASR subunits. Thus, as observed previously, the majority of nuclear Rbfox1 is associated stoichiometrically with LASR on chromatin (Damianov et al. 2016; Jiang et al. 2021; Ying et al. 2017).

**Figure 1.**
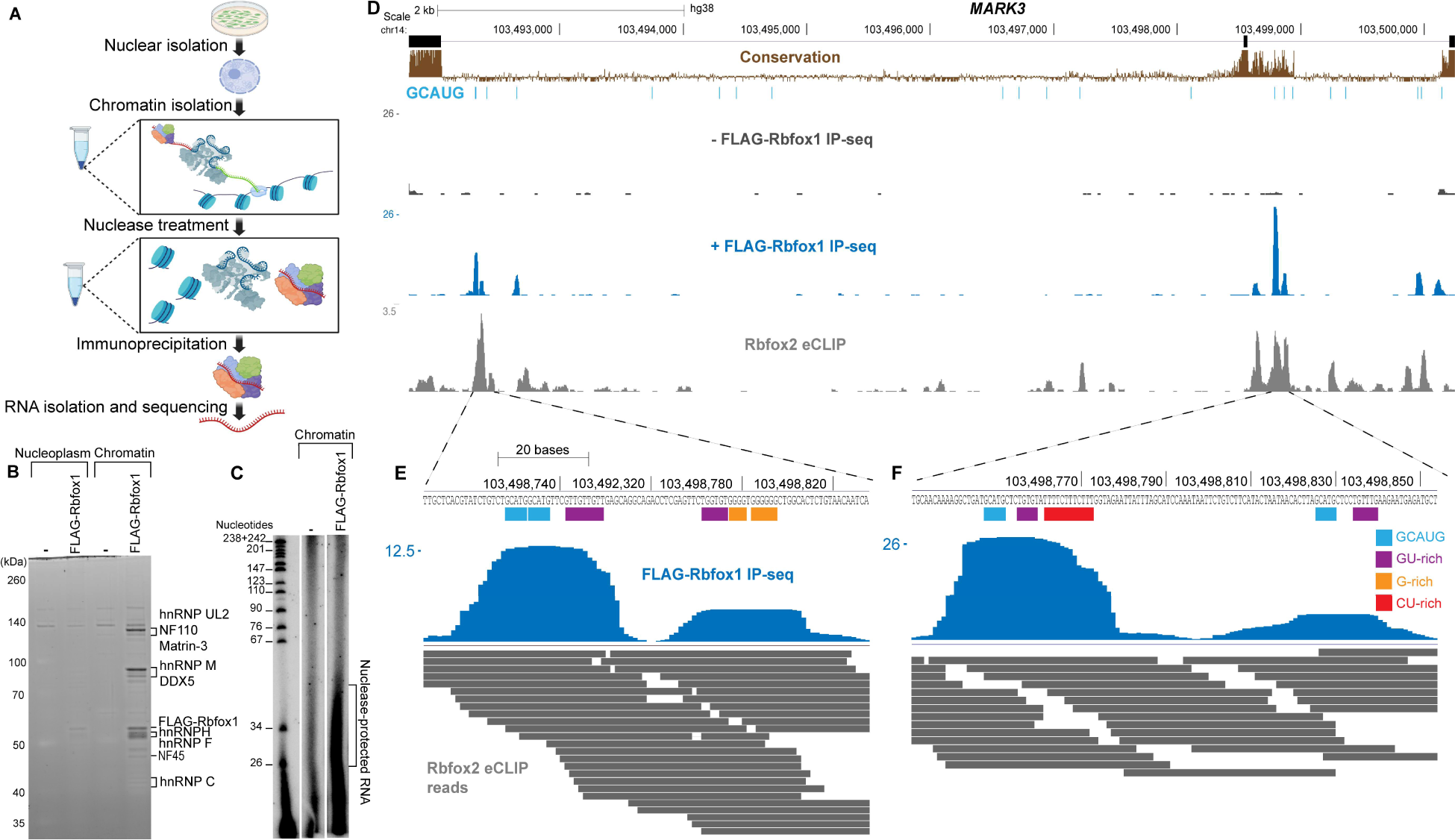
Mapping Rbfox1/LASR binding sites on chromatin-associated RNA by nuclease-protection. A) Diagram of IP-seq method for purifying nuclease-protected RNA associated with chromatin-enriched protein complexes. B) SYPRO Ruby stained SDS-PAGE of immunoprecipitations from parental (-) and FLAG-Rbfox1 expressing HEK293 cells. C) Urea-PAGE of P32 labeled nuclease-protected RNAs isolated with FLAG-Rbfox1. D) UCSC Genome Browser tracks of nuclease-protected fragments in the introns flanking *MARK3* exon 16. Light blue bars indicate position of GCAUG elements and the phyloP conservation across 100 vertebrae species is displayed in brown. The IP-seq browser track in black is from parental cells (-) and the blue is from FLAG tagged Rbfox1 expressing cells. Rbfox2 eCLIP from HEK293T cells from Van Nostrand et al. is shown in gray. E-F) Two regions containing protected sites are expanded to display their nucleotide composition. Motifs present within these regions are delineated by colored bars as the following: GCAUG (blue), GU-rich (purple), G-rich (orange), and CU-rich (red). Individual reads in this region from the Rbfox2 eCLIP are displayed in grey.

To examine whether RNA co-purified with the complex, the immunoprecipitated material was deproteinized, extracted with phenol-chloroform, DNAse treated, dephosphorylated, and 5’ end-labeled with [γ-^32^P]ATP. This material was then analyzed by Urea-PAGE and autoradiography. As shown in Figure 1C, small RNA fragments less than 50 nucleotides long copurified with Rbfox1/LASR, presumably protected from nuclease degradation by their interaction with the complex. To identify these RNAs, we converted fragments of 20-50 nucleotides into cDNA using a modified iCLIP library protocol and sequenced them.

The FLAG-Rbfox1 IP-seq yielded 16.5 to 32.7 million reads per replicate with an average length of 32 nucleotides (Fig. S1A). Across the three replicates, greater than 60% of the reads mapped to unique locations in the genome, forming clusters of aligned reads. We defined 561,273 clusters that contained at least 10 reads in merged replicates using the peak caller YODEL (Palmer et al. 2017). Control IP-seq from cells that do not express FLAG-Rbfox1 yielded 14 million reads with a 68% mapping rate. Interestingly, many of the reads from the control cells aligned at intron branchpoints, similar to what was previously observed with isolated U2 snRNPs (Fig. S1B). These reads from the control cells are attributed to weak cross-reactivity of the anti-FLAG antibody with a subunit of the U2 snRNP. To distinguish bona fide binding sites from background, we filtered for clusters that were enriched in the experimental set over the background set, leaving 472,757 clusters. These clusters were mostly intragenic, with the majority mapping to introns or 3’ UTRs, similar to Rbfox binding patterns measured by CLIP (Fig. S1C-D).

We then assessed the overlap between Rbfox1/LASR IP-seq clusters and a previously reported set of Rbfox2 eCLIP clusters from HEK293 cells (Van Nostrand et al. 2016). In previous CLIP studies, binding sites of all three Rbfox proteins largely overlapped (Damianov et al. 2016; Weyn-Vanhentenryck et al. 2014). We thus expected the patterns of binding of Rbfox1 by IP-seq and Rbfox2 by eCLIP to be similar. The IP-seq and eCLIP clusters were each segregated into two groups based on the presence or absence of a GCAUG element. Approximately 50% of GCAUG-containing Rbfox2 eCLIP clusters overlapped with GCAUG-containing IP-seq clusters (Fig. S2A). Beside differences in Rbfox1 and Rbfox2 targeting, several factors could limit the overlap between the eCLIP and IP-seq clusters. eCLIP and IP-seq clusters were defined using different computational tools and statistical assessments that likely result in differences in the sensitivity and specificity of cluster detection. Each method may also fail to report particular interactions. For example, some Rbfox binding sites might not crosslink and therefore be undetected by eCLIP, whereas some interactions may not be stable during the nuclease treatment and be undetected by IP-seq. Both methods also generated substantial numbers of clusters that lack GCAUG. Comparing these groups, approximately 18% of Rbfox2 eCLIP clusters without GCAUG overlapped with IP-seq clusters lacking this motif (Fig. S2B). The more limited overlap of clusters without GCAUG is expected since eCLIP will mostly detect direct Rbfox binding sites whereas IP-seq will also detect sites protected by LASR.

Figure 1D displays a genome browser view of IP-seq protected regions that map in introns adjacent to an alternative exon of *MARK3*. The patterns of protected fragments isolated by IP-seq and of crosslinking sites generated by eCLIP are very similar. Two peaks of protected fragments are adjacent to the 5’ splice site of the upstream intron (Fig. 1E). These peaks are more tightly distributed than the reads of CLIP crosslink sites. The two IP-seq clusters in this region contain distinct sets of motifs. The first cluster contains two GCAUG elements along with a downstream GU-rich element. The second cluster lacks GCAUG motifs and instead contains G-rich and GU-rich elements. These non-GCAUG motifs within the protected fragments are potential binding sites for the LASR subunits hnRNP M, which binds GU-rich motifs (Feng et al. 2019; Ho et al. 2021; Zhu et al. 2022), and hnRNP H/F, which bind G-rich elements (Caputi and Zahler 2001; Dominguez et al. 2010; Dominguez and Allain 2006; Matunis et al. 1994; Penumutchu et al. 2018; Uren et al. 2016; Van Nostrand et al. 2020). The eCLIP reads in this region are distributed across all these elements. A similar pattern is observed in the IP-seq protected fragments adjacent to the 5’ splice site of the downstream intron (Fig. 1F). In this region the protected fragments align in two peaks, and each contains a GCAUG element plus other motifs. The eCLIP reads are again more broadly distributed across this region. Rbfox crosslinking to regions without GCAUG may be due to its proximity after recruitment to the RNA as part of the LASR complex, whose subunits can bind to these sites.

### RNA protected by Rbfox1/LASR is enriched in binding motifs for both Rbfox and LASR subunits

We next identified RNA motifs that were enriched in IP-seq isolated intronic fragments relative to their frequency in their total intron sequence. HOMER analysis revealed the canonical Rbfox UGCAUG element as the most enriched motif, found in 25% of the reads (Fig. 2A) (Heinz et al. 2010). Additionally, 30% of the reads contain the GCAUG motif, while secondary Rbfox motifs (e.g. GCACG) were less prevalent though still significant (Fig. S3). Other motifs enriched in the protected fragments matched those defined for individual LASR subunits. GU-rich elements (GUGUGU, GUUGUU) that are known to bind hnRNP M were present in at least 21% of reads, nearly as enriched as Rbfox motifs. The GUGUGU motif has also been reported to be a secondary Rbfox motif (Begg et al. 2020). However, its greater abundance in the protected fragments compared to other secondary Rbfox motifs suggests that its isolation is likely due to a LASR subunit interaction rather than Rbfox. G-rich, U-rich, and CU-rich elements, motifs previously reported to bind to hnRNP H/F, hnRNP C, and Matrin3 respectively, were each present in at least 10% of reads (Ramesh et al. 2020; Uemura et al. 2017; Zarnack et al. 2013). Also among the enriched sequences identified by HOMER was a motif containing UAG whose cognate binding factor is not yet clear.

**Figure 2.**
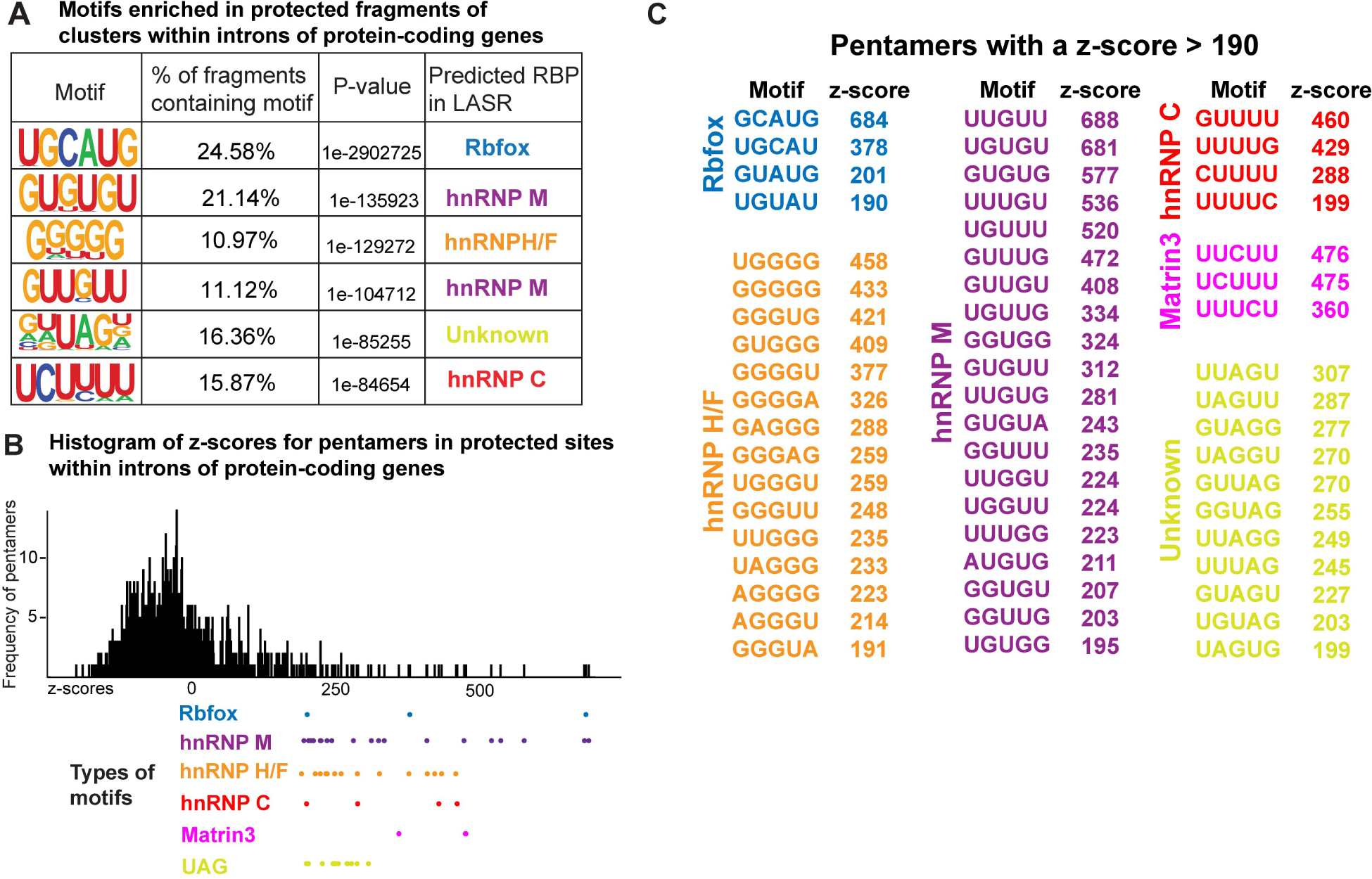
RNA protected by Rbfox1/LASR is enriched in binding motifs for both Rbfox and LASR subunits. A) HOMER analysis of enriched motifs of 4-6 nucleotides in protected fragments within clusters in introns of protein-coding genes. The enrichment of motifs was determined by comparing their frequency in protected fragments to randomly sampled regions from introns that contained at least one IP-seq cluster. B) Z-score analysis of pentamers within clusters in introns of protein-coding genes. C) Pentamers with a z-score of more than 190, which includes motifs that have a z-score in the top 5% of the distribution, are categorized and displayed.

In an alternative approach to HOMER, we compared the frequency of pentamers within the intronic IP-seq clusters with their frequencies in randomly sampled regions from the same introns. Z-scores for these pentamers were calculated, and their distribution is illustrated in Figure 2B. Pentamers with z-scores above 190, which include the top 5% of the pentamers within the population, are highlighted and their sequences are shown in Figure 2C. All these pentamers can be classified into motif categories identified in the HOMER analysis. The motifs clustered together with the highest z-scores are GCAUG (z=684), UUGUU (z=688), and UGUGU (z=681). A population of GU-rich, G-rich, CU-rich, and U-rich elements have z-scores between 300-600 and another population have z-scores below 300. The most enriched UAG containing element, UUAGU, has a z-score of 307. Z-score analysis of hexamers gave similar results (Supplementary Table 1).

### RNA bound to Rbfox1/LASR contains modules of multi-part motifs

We found that 13% of GCAUG pentamer motifs and 27% of the UGCAUG hexamers within expressed transcripts were recovered in protected RNA fragments (Fig. S5A). This observation is consistent with CLIP studies where not all GCAUG elements crosslink to Rbfox (Damianov et al. 2016; Jangi et al. 2014; Lovci et al. 2013; Weyn-Vanhentenryck et al. 2014). Individual RBPs within some RNPs can each directly contact portions of contiguous motifs along an RNA strand (Hennig et al. 2014; Kuwasako et al. 2014; Wysoczański et al. 2014). Therefore, we examined co-occurrence of motifs within the same protected fragments. Combined recognition of such elements may increase binding affinity, and the absence of gaps between the elements may confer greater resistance to nuclease digestion. Approximately 90% of protected fragments with GCAUG contained at least one other enriched binding motif, with U-rich elements being the most common, appearing in 66% of GCAUG-containing fragments (Fig. 3A). GU-rich elements were also common, found in 52% of GCAUG-protected fragments. Binding of multiple proteins to the same RNA fragment might also extend the length of the nuclease-protected region. Accordingly, protected fragments containing GCAUG alongside another enriched motif exhibited a mean length of 34 nucleotides, compared to 28 nucleotides for fragments containing only GCAUG and no other enriched motifs (Fig. 3B). The presence of different types of motifs in reads is also correlated with different distributions of read lengths (Fig. S4). These results suggest that Rbfox and LASR can have multiple contact points within the same segment of RNA. Examples of protected fragments composed of multiple motifs are shown for the *MAD1L1* and *LAMP2* transcripts in Figure 3C.

**Figure 3.**
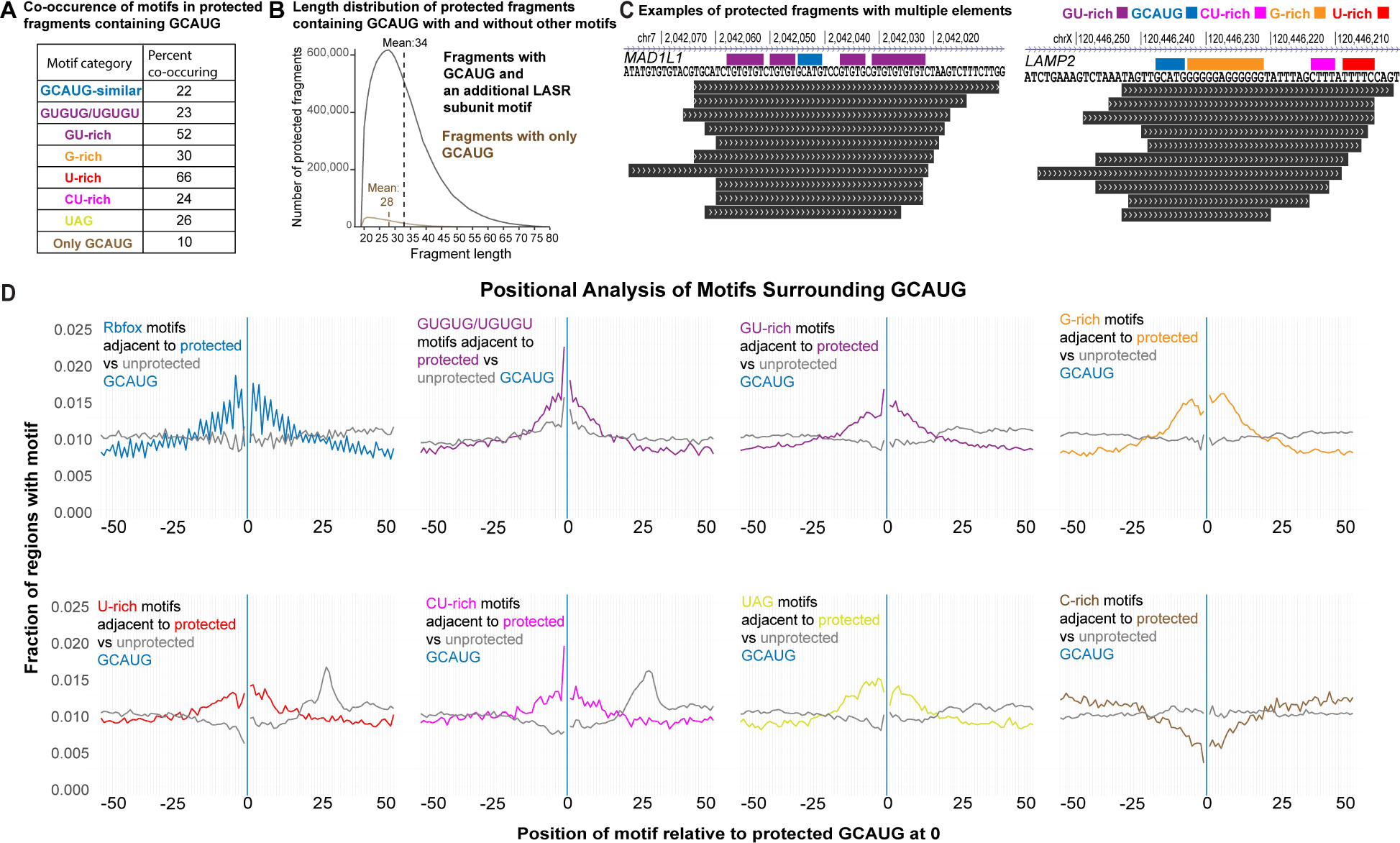
RNA bound to Rbfox1/LASR contains modules of multi-part motifs. A) Percent of protected fragments that contain GCAUG and other specified motifs. GCAUG-similar motifs (blue) include all Rbfox secondary motifs shown in Supplementary Figure 2. GU-rich (purple) motifs consist of all pentamers under hnRNP M in Figure 2, except for GUGUG and UGUGU, which form their own category. U-rich (red) motifs cover all pentamers associated with hnRNP C, CU-rich (pink) includes all motifs under Matrin3, and UAG (yellow), encompasses all motifs labeled unknown. B) Distribution of length of protected fragments that contain GCAUG and motifs of LASR subunits (black) compared to fragments with GCAUG that do not contain these motifs (brown). Dotted lines are at the mean of the distribution. C) Examples of protected fragments containing combinations of motifs in the *MAD1L1* and *LAMP2* transcripts. Motifs are outlined by colored bars that correspond to the same color scheme in *A*. D) Positional analysis of motifs surrounding GCAUG. Graphs display the fraction of regions containing each type of motif relative to GCAUG at position 0, indicated by a vertical blue line. A region extending 50 nucleotides upstream and downstream of the GCAUG was defined as the surrounding region. The frequency of motifs surrounding protected GCAUGs is plotted with the same colors defined in *A*, while the frequency of motifs surrounding unprotected GCAUGs is plotted in grey. Motif categories follow the same descriptions as *A* except for Rbfox motifs which include GCAUG in addition to the Rbfox secondary motifs. C-rich motifs (brown) include ten pentamers with the lowest z-scores: AAACA, CCAGG, GCCAC, CCUCA, GCCCA, CAGCC, CUCCC, CCAGC, UCCCA, and CCCAG.

To examine the placement of motifs relative to the GCAUG, we plotted their frequency along sequences adjacent to protected GCAUG motifs (Fig. 3D). LASR motifs were most common immediately adjacent to the protected GCAUG, within 1-10 nucleotides upstream and downstream. GU-rich, and CU-rich motifs were particularly common upstream of the GCAUG, presumably reflecting the preference for U as the initial nucleotide of UGCAUG. These motifs usually abut the GCAUG, while G-rich motifs are more frequently spaced 5-6 nucleotides away, suggesting a possible structural constraint on the co-binding of the hnRNP H/F and Rbfox proteins. As a control, we assessed the occurrence of these motifs adjacent to unprotected GCAUGs and found that none of the motifs were enriched nearby (Fig. 3D). Notably, U-rich and CU-rich motifs showed a high occurrence approximately 25 nucleotides downstream of unprotected GCAUGs, indicating that a motif at this distance may disfavor binding of Rbfox to the GCAUG. We also assessed the occurrence of C-rich motifs, which are not enriched in the Rbfox1/LASR protected RNA, and found that these elements are depleted in sequences adjacent to protected GCAUGs. In addition to examining the positional frequency of LASR motifs in genomic regions adjacent to protected GCAUGs, we also analyzed their frequency within protected fragments that contained a GCAUG. This analysis shows similar patterns of positional motif enrichment relative to the GCAUG (Fig. S5B).

### Rbfox1/LASR containing the F125A RNA binding mutant of Rbfox1 loses binding to GCAUG but not the LASR elements

To examine the contribution of the Rbfox RRM to the isolation of the protected fragments, we analyzed a mutant Rbfox1 protein, Rbfox1(F125A). This mutation eliminates a critical phenylalanine on the RNA binding surface of the RRM domain, increasing the K_d_ of the UGCAUG interaction approximately 1500 fold (Auweter et al. 2006). A FLAG-Rbfox1(F125A) transgene was integrated into the Rbfox2^−/−^ cells and the protein was isolated via its epitope tag. Total protein staining of the immunoprecipitates yielded a banding pattern matching that of wildtype FLAG-Rbfox1, indicating the mutant and wildtype Rbfox1 both interact with LASR (Fig. 4A). RNA isolated from the Rbfox1(F125A)/LASR complex had a similar Urea-PAGE profile to the RNA protected by Rbfox1(WT)/LASR (Fig. 4B). This protected RNA was sequenced and had similar average lengths and mapping rates to the wildtype IP-seq (Fig. S1A).

**Figure 4.**
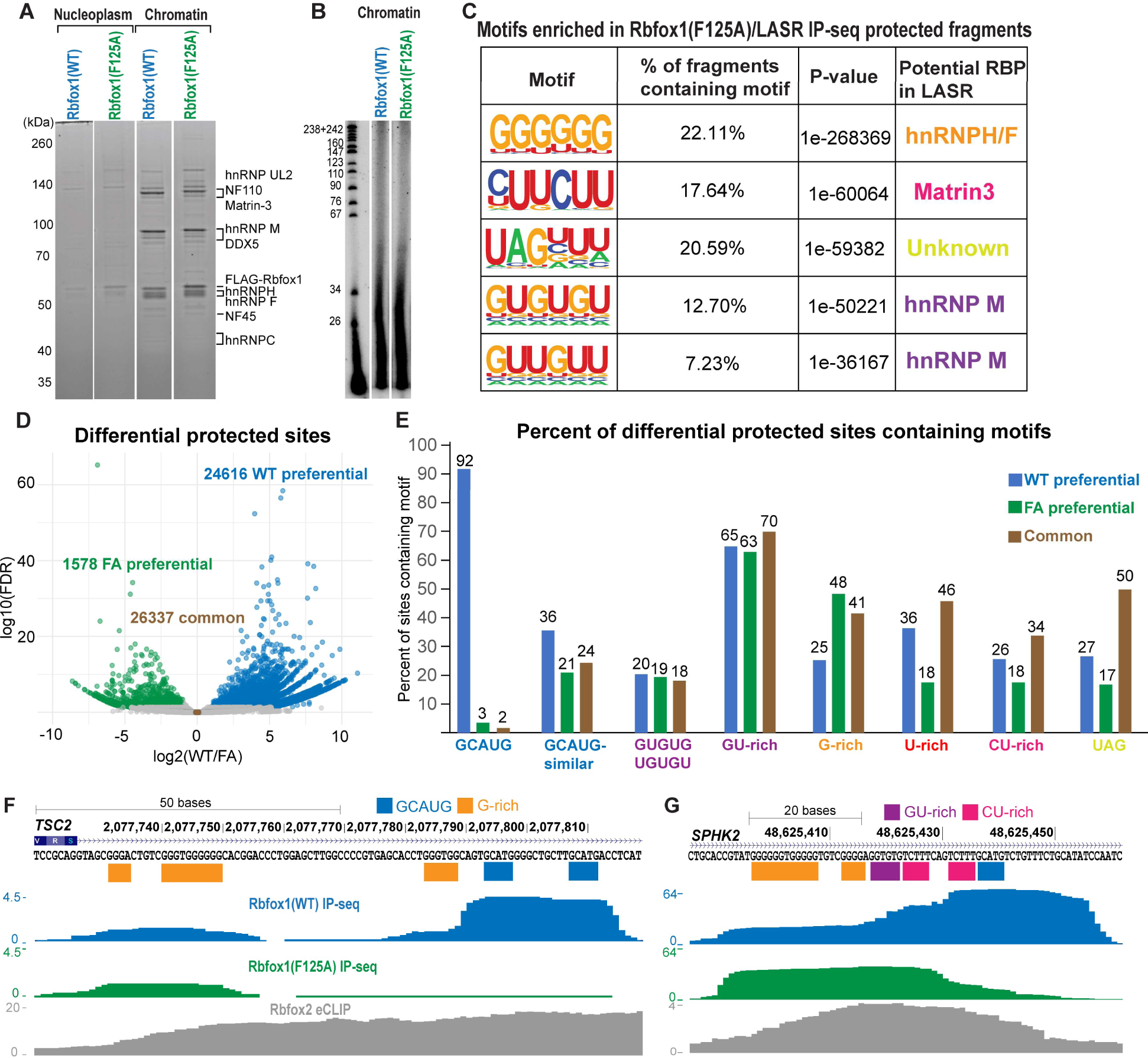
Rbfox1/LASR containing the F125A RNA binding mutant of Rbfox1 loses binding to GCAUG but not the LASR elements. A) SYPRO Ruby stained SDS-PAGE of immunoprecipitation from FLAG-Rbfox1(WT) and FLAG-Rbfox1(F125A) expressing cells. B) Urea-PAGE of P32 labeled nuclease-protected RNAs purified from immunoprecipitations of FLAG-Rbfox1(WT) and FLAG-Rbfox1(F125A). C) Enriched motifs identified by HOMER in protected fragments of Rbfox1(F125A)/LASR from clusters in introns of protein-coding genes. D) Volcano plot illustrating differential binding in Rbfox1(WT)/LASR versus Rbfox1(F125A)/LASR as analyzed by DESeq2. Sites preferentially bound in the WT IP-seq are shown in blue, FA preferential sites are shown in green, and common sites are displayed in brown. E) Analysis of occurrence of motifs in differentially bound sites with the same color schemes as *D*. F-G) UCSC Genome Browser view of nuclease-protected fragments within the intron downstream of *TSC2* exon 25 (F) and within an intronic region in *SPHK2* (G). Positions of enriched motifs in protected sites are indicated with colored bars as the following: GCAUG (blue), G-rich (orange), GU-rich (purple), and CU-rich (pink).

RNA fragments isolated with Rbfox1(F125A)/LASR exhibited different motif frequencies than Rbfox1(WT) (Fig. 4C). The GCAUG motif was no longer enriched, while all the other motifs enriched with Rbfox1(WT)/LASR complex remained, although their frequencies shifted (Fig. 4C, Fig. S6). G-rich (hnRNP H/F) and CU-rich elements (Matrin3) were more frequent compared to wildtype. GU-rich elements (hnRNP M sites) decreased in frequency when Rbfox binding was lost. DESeq2 was used to analyze differential binding between the LASR complexes containing wildtype and F125A Rbfox1 (Fig. 4D) (Love et al. 2014). We classified 24,616 sites as WT preferential, characterized by a log2(WT/FA) > 1 and an FDR < 0.05. A total of 26,337 sites were defined as unchanged and therefore common between WT and the FA mutant, with log2(WT/FA) values ranging between −0.1 and 0.1. Fewer sites (1,578) were identified as FA preferential, with log2(WT/FA) < −1 and an FDR < 0.05. The remaining sites, with log2(WT/FA) values between −0.1 to −1 and 0.1 to 1, were unclassified.

We then analyzed motif prevalence in differentially or commonly bound sites (Fig. 4E). GCAUG appeared in 92% of WT preferential sites, but only 3% of FA preferential and 2% of commonly bound sites. Rbfox secondary motifs (GCUUG, GAAUG, GCACG, GUAUG, GUUUG) were moderately more prevalent in WT preferential sites (32%) compared to FA preferential (21%) and commonly bound (24%) sites. Conversely, G-rich elements were more frequent in the common sites (41%) and FA preferential sites (48%) than in WT preferential sites (25%). GU-rich and GUGUG/UGUGU motifs were similarly prevalent across all sites, 63-70% and 18-20% respectively. U-rich, CU-rich, or UAG containing elements all had the highest prevalence in commonly bound sites, followed by WT preferential, and the lowest in FA preferential sites.

An example of these differentially protected sites is downstream of an Rbfox-regulated exon in the *TSC2* transcript, a gene essential for cell growth (Huang et al. 2008) (Fig. 4F). eCLIP of Rbfox2 exhibited broadly distributed crosslinking across the entire region downstream of the exon (Van Nostrand et al. 2016). In contrast, IP-seq produced two clear peaks of protected fragments. The more proximal peak, containing several conserved G-rich elements, was bound by both wildtype Rbfox1 and the F125A mutant, indicating that these elements are protected independently of Rbfox’s RRM, most likely by hnRNP H/F. A second downstream cluster containing conserved GCAUG motifs was isolated with the wildtype Rbfox1 but disappeared in the F125A mutant. Thus, Rbfox’s selective affinity for GCAUG is essential for binding to the downstream region and protecting it from nuclease cleavage.

In the *SPHK2* transcript, differential protection by Rbfox1 versus LASR was observed within a single protected region (Fig. 4G). In the Rbfox1(WT) IP-seq, the predominant peak aligns with a single UGCAUG, but extends upstream to include GU-rich, CU-rich, and G-rich elements. In the Rbfox1(F125A) IP-seq, protection at the UGCAUG site and the upstream CU-rich site decreases significantly, while protection of the G-rich and GU-rich regions is maintained. Comparison of IP-seq between wildtype and F125A Rbfox1 can therefore identify Rbfox1-bound GCAUG elements even when they are directly adjacent to other elements. We also observed increased binding to certain sites by the Rbfox(F125A)/LASR. Examples of such sites in the *MSPP2* and *ARFGAP1* transcripts are shown in Supplementary Figure 7. Consistent with the motif distribution analysis, these sites are composed of a combination of LASR elements and lack GCAUG.

### Rbfox1(WT)/LASR and Rbfox1(F125A)/LASR regulate splicing through distinct binding sites adjacent to cassette exons

Given that LASR in complex with Rbfox1(F125A) continues to bind numerous sites, including some preferentially over wildtype, we assessed its ability to regulate splicing. Splicing in cells lacking Rbfox was compared with cells expressing comparable levels of either wildtype Rbfox1 or the F125A mutant (Fig. 5A). PolyA(+) RNA was isolated and subjected to paired-end RNA-seq. rMATS-turbo was used to identify splicing changes between the three conditions (Wang et al. 2024). Each condition exhibited a distinct pattern of splicing regulation (Fig. 5B, Fig. S8). 462 exons were uniquely regulated by the wildtype Rbfox1 (WT-regulated) whereas 542 exons were affected by both the wildtype and F125A mutant (WT/FA-regulated). A third set of exons, 289, were altered by the F125A mutant but not by the wildtype (FA-regulated).

**Figure 5.**
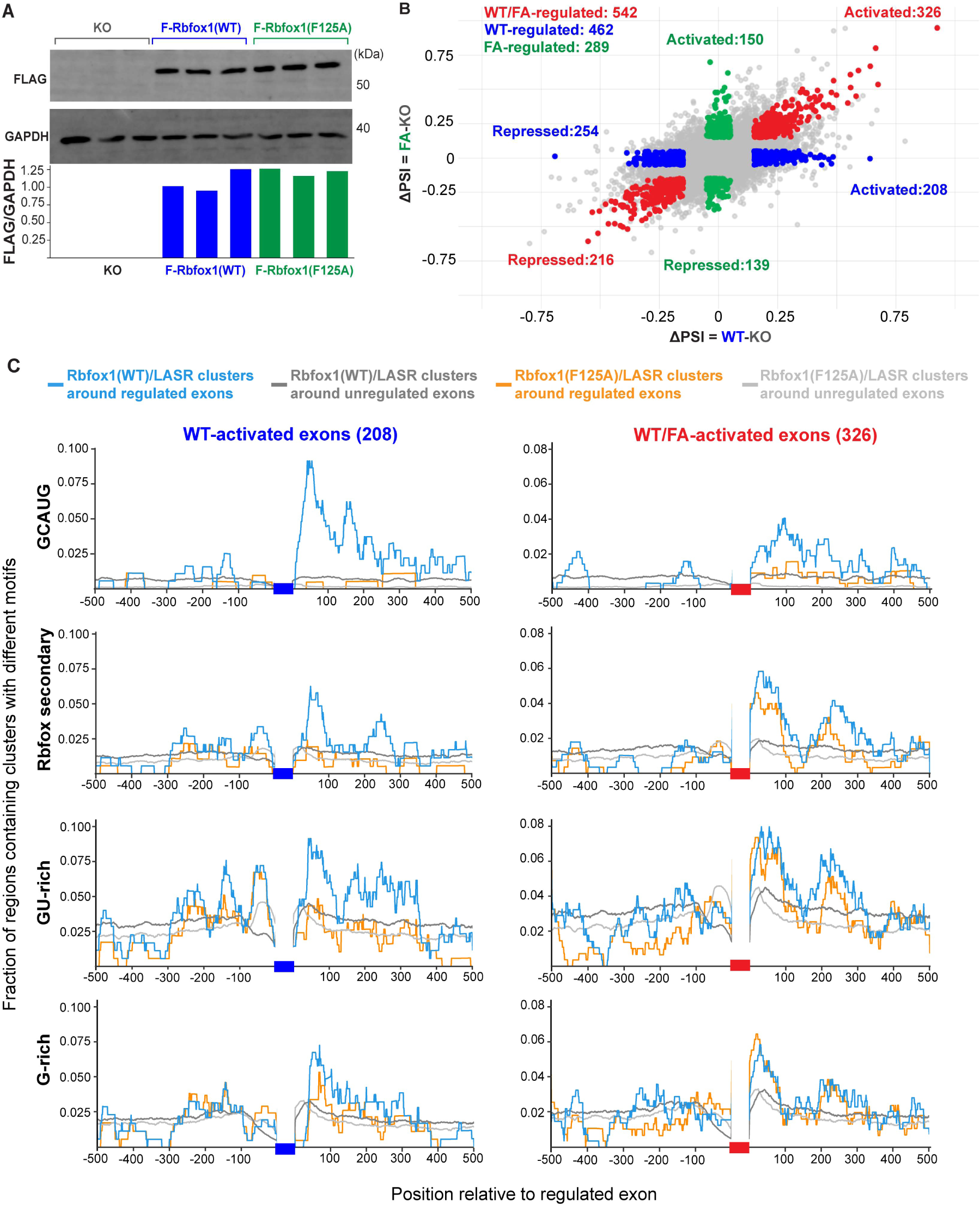
Rbfox1(WT)/LASR and Rbfox1(F125A)/LASR regulate splicing through distinct binding sites adjacent to cassette exons. A) Immunoblot of FLAG and GAPDH in HEK293 Rbfox deficient, FLAG-Rbfox1(WT), and FLAG-Rbfox1(F125A) expressing cells, B) Comparison of regulated exons in Rbfox1(WT) and Rbfox1(F125A) expressing cells. The x-axis is the ΔPSI calculated from subtracting PSI of the exon in Rbfox1(WT) cells from PSI of the exon in Rbfox KO cells. The y-axis is the ΔPSI from subtracting PSI of the exon in Rbfox1(F125A) cells from PSI of exons in Rbfox KO cells. All exons in red, green, and blue have an FDR < 0.05. Exons with |ΔPSI(WT-KO)| and |ΔPSI(FA-KO)| >= 0.15 where |ΔPSI(WT-KO) - ΔPSI(FA-KO)| <= 0.15 are considered regulated by both WT and FA and are colored red; exons that have a |ΔPSI(WT-KO)| >= 0.15 and |ΔPSI(FA-KO)| <= 0.05 are considered WT-regulated and are colored blue; exons that have a |ΔPSI(FA-KO)| >= 0.15 and |ΔPSI (WT-KO)| <= 0.05 are considered FA-regulated and are colored green. All other exons are colored grey. C) RNA binding map of Rbfox1(WT)/LASR and Rbf1ox1(F125A) protected sites 500 bp upstream and downstream of WT-activated, WT/FA-activated, FA-activated exons, and unregulated exons. Unregulated exons are those with |ΔPSI(WT-KO)| and |ΔPSI(FA-KO)| <= 0.01. Clusters are categorized based on the motifs they contain: GCAUG, Rbfox secondary (including GCUUG, GAAUG, GCACG, GUAUG, GUUUG), GU-rich, and G-rich motifs. For regulated exons, the frequency of Rbfox1(WT)/LASR clusters is plotted in light blue while the Rbfox1(F125A)/LASR clusters are displayed in orange. The frequency of Rbfox1(WT)/LASR clusters around unregulated exons is plotted in dark grey and the frequency of Rbfox1(F125A)/LASR clusters is light grey.

RBPs often have position-dependent effects on their regulatory targets, with Rbfox typically activating exons when bound downstream and inhibiting them when bound upstream (Conboy 2017). To examine how protected regions affected splicing, we analyzed the frequencies of IP-seq clusters containing different motifs within 500 nucleotides upstream and downstream of exons in each regulatory class (Fig. 5C, Fig. S9). As a control, we also assessed the frequency of these clusters around exons unaffected by either the either wildtype or F125A mutant Rbfox1. Clusters from Rbfox1(WT) and Rbfox1(F125A) IP-seq were categorized based on the presence of particular motifs including: GCAUG, Rbfox-secondary (GCUUG, GAAUG, GCACG, GUAUG, GUUUG), GU-rich (divided to GUGUG/UGUGU and other GU-rich pentamers), G-rich, CU-rich, and U-rich elements. The enrichment of these clusters around regulated exons, compared to unregulated exons, was assessed via a bootstrapping test (Supplementary Tables 2-7).

‘For exons activated exclusively by wildtype Rbfox1 (Fig. 5C, first column), Rbfox1(WT)/LASR IP-seq clusters containing the GCAUG motif were enriched downstream, consistent with previous results indicating that downstream binding leads to splicing activation. In contrast, Rbfox1(F125A)/LASR clusters with GCAUG were seldom found near the exons activated by the wildtype Rbfox1. Rbfox1(WT)/LASR clusters containing Rbfox secondary motifs, GU-rich, G-rich, U-rich, and CU-rich elements were also more frequent downstream of WT-activated exons (Fig. 5C, Fig. S9B). These data indicate that the activation of exons by the wildtype Rbfox1 is dependent on binding to downstream GCAUG, Rbfox secondary, and LASR RNA elements.

In contrast to WT-activated exons, the WT/FA-activated set showed a more limited downstream enrichment of Rbfox1(WT)/LASR clusters with GCAUG (Fig. 5C, second column). Interestingly, both Rbfox1(WT)/LASR and Rbfox1(F125A)/LASR clusters containing binding elements for LASR were enriched downstream of the WT/FA-activated exons (compare blue and yellow lines in Fig. 5C and S9B, second column). This binding profile suggests that these exons are activated by binding of LASR to its regulatory elements, whether or not the Rbfox RRM can engage GCAUG. There was also a similar occurrence of Rbfox1(WT)/LASR and Rbfox1(FA)/LASR IP-seq clusters containing Rbfox secondary motifs downstream of these exons, indicating Rbfox may normally contact these secondary motifs but LASR binding makes this less essential than for exons controlled by higher affinity Rbfox primary motifs.

Exons activated only by the F125A mutant protein and their surrounding clusters were fewer in number than the other two groups, making enrichments difficult to discern. Rbfox1(WT)/LASR and Rbfox1(F125A)/LASR clusters containing GCAUG showed no obvious positional bias on these exons (Fig. S9A). Interestingly, there were enrichments upstream of some of these exons for Rbfox1(WT)/LASR and Rbfox1(F125A)/LASR clusters containing Rbfox secondary, GU-rich, and G-rich motifs. These binding patterns raise the possibility that the F125A mutant may be able to act as a dominant negative factor to reverse splicing repression by certain factors. Such a dominant negative effect has been observed by a splice variant of Rbfox that lacks a portion of the RNA-binding domain (Damianov and Black 2010; Nutter et al. 2016).

We also assessed binding of Rbfox1/LASR around repressed exons (Fig. S9C). For exons repressed only by the wildtype Rbfox1, the Rbfox1(WT)/LASR clusters containing GCAUG were moderately enriched upstream compared to the Rbfox1(F125A)/LASR clusters. This position-dependent effect on splicing repression is similar to the pattern observed in CLIP studies, but less pronounced (Jangi et al. 2014; Lovci et al. 2013; Moss et al. 2023; Weyn-Vanhentenryck et al. 2014; Zhou et al. 2021). The occurrence of protected sites containing Rbfox secondary, GU-rich, and G-rich elements did differ dramatically between wildtype and F125A IP-seq for the WT-repressed exons. In the WT/FA-repressed and FA-repressed exons, the prevalence of protected sites containing any of the motifs was low in both the wildtype and the F125A IP-seq, with no clear patterns of enrichment. The mechanism of exon repression by Rbfox1/LASR appears more complex than the binding that determines exon inclusion.

### Multipart elements within Rbfox1/LASR binding sites have combined effects on exon inclusion

We next tested the regulatory effects of individual motifs within protected sites adjacent to regulated exons. Exon 16 of the *CAMKK2* transcript was previously reported to be regulated by PKA signaling and involved in neurite branching (Cao et al. 2011). This exon is strongly activated by wildtype Rbfox1 and not the F125A mutant (Fig. 6A), within a complex set of alternative splicing changes that include stimulation of a new downstream 3’ exon. The intron downstream of exon 16 contains three IP-seq clusters (Fig. 6A). The most proximal cluster, region 1, contains G-rich elements, and a more distal region 2 contains two GCAUG motifs surrounding a G-rich element. A third protected segment, region 3, contains a GCAUG motif. All three protected regions were isolated with Rbfox1(WT)/LASR; but binding to regions 1 and 2 was greatly reduced, and binding to region 3 eliminated by the F125A mutation.

**Figure 6.**
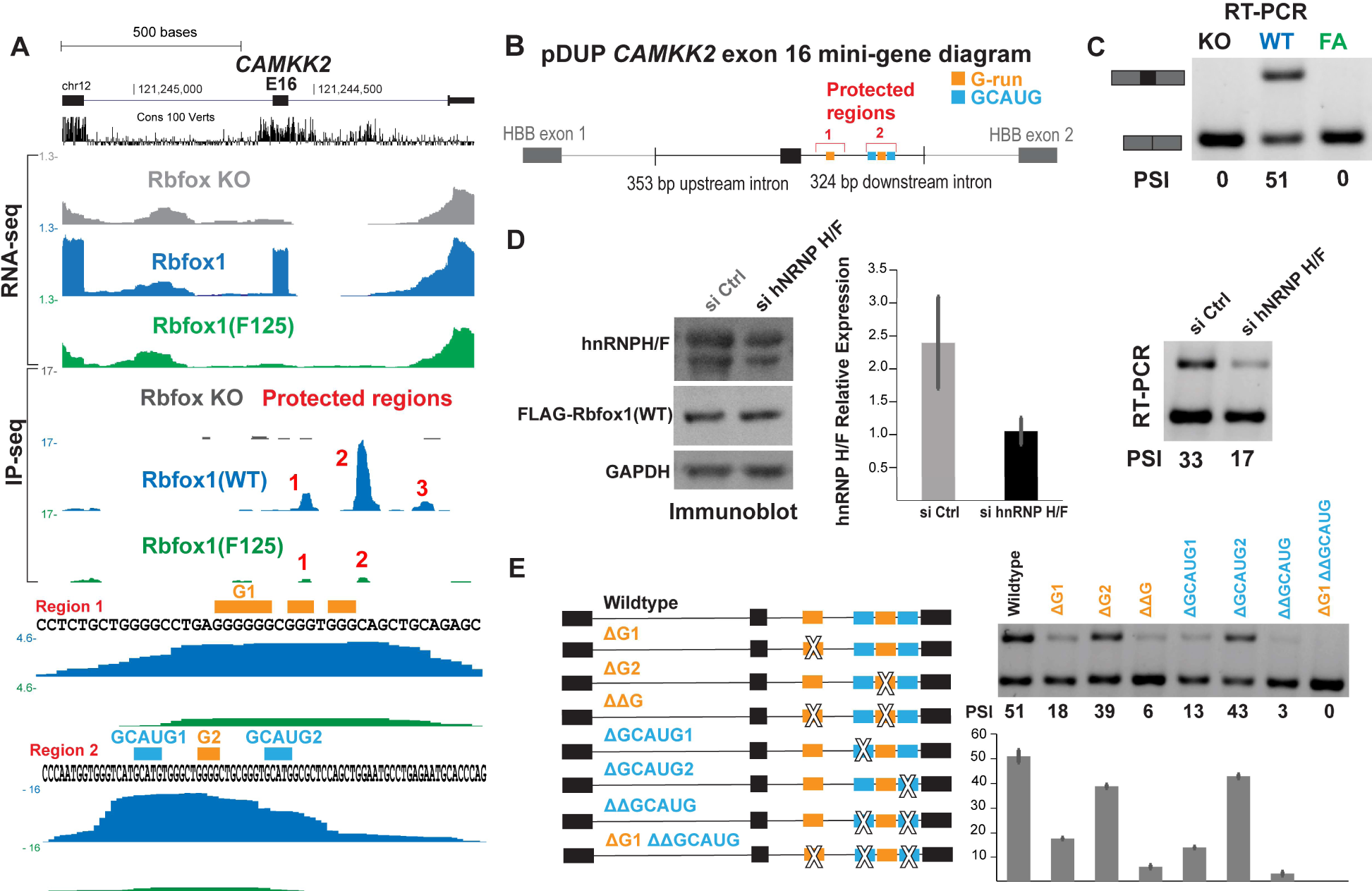
Multipart elements within Rbfox1/LASR binding sites have combined effects on exon inclusion. A) Genome browser view of *CAMKK2* transcript with RNA-seq tracks from Rbfox deficient (grey), Rbfox1(WT) (blue), and Rbfox1(F125A) (green) expressing cells. A track containing conservation across vertebrates is displayed in black. IP-seq tracks, displayed below the RNA-seq, show three protected regions downstream of a Rbfox1(WT) activated exons outlined with red text. The sequence features of two of the protected regions are displayed in more detail at the bottom. Motifs are outlined by colored bars with G-rich elements in orange and GCAUG in light blue. B) Diagram of the *CAMKK2* exon 16 DUP mini-gene. The *CAMKK2* fragment is inserted between *HBB* exons one and two and their intervening intron. Protected regions and the elements within them are outlined per the same color scheme as *A.* C) Agarose gel electrophoresis of the CAMKK2 exon 16 DUP mini-gene splice products amplified by RT-PCR. D) Immunoblot of hnRNPH/F siRNA knockdown and RT-PCR splicing assay of the *CAMKK2* exon 16 DUP-mini gene in these conditions. E) Diagram of deletion mutants of *CAMKK2* exon 16 DUP mini-gene along with RT-PCR analysis of mutants.

To assess the regulatory effects of these regions, we adapted a mini-gene construct containing *CAMKK2* exon 16, along with 353 nucleotides of upstream and 324 nucleotides of downstream intronic sequence (Cao et al. 2011). This fragment contained the first two protected regions isolated with Rbfox1/LASR and lacks the third region. We integrated this fragment into a DUP backbone where all GCAUG elements were eliminated to avoid their possible confounding regulatory effects (Fig. 6B). This mini-gene was transiently expressed in the Rbfox^−/−^, Rbfox1(WT), and Rbfox1(F125A) cell lines, with its spliced products assayed by RT-PCR. The splicing of this mini-gene recapitulated that of the endogenous gene, with exon 16 completely skipped from the mRNA in the Rbfox2^−/−^ and Rbfox1(F125A) cells, but strongly included in the Rbfox1(WT) cells (Percent spliced in, PSI = 51%, Fig. 6C).

Given that hnRNP H and F can bind to G-rich elements, we assessed splicing after siRNA-mediated depletion of these proteins. The siRNA treatment of the Rbfox1(WT) cells reduced hnRNP H/F protein expression by about 70% (Fig. 6D). The exon PSI in the control siRNA treatment was 33% compared to 17% after the hnRNPH/F knockdown. Thus, *CAMKK2* exon 16 is highly sensitive to hnRNP/F, which stimulate its splicing.

To assess the regulatory contributions of individual motifs present in the two protected regions, we constructed a series of deletion mutants (Fig. 6E). G-rich elements were removed from the first or second protected regions, or from both. Loss of the first G-rich element reduced exon 16 splicing from 51% to 18%, while loss of the second G-rich element reduced splicing to 39%. Double mutation of both G-rich elements further reduced the PSI to 6%. These two G-rich elements are thus positive regulators of exon 16 and have additive effects on its splicing.

The GCAUG elements also had effects on exon inclusion. Deletion of the first GCAUG motif decreased PSI from 51% to 13%, while excising the second reduced PSI to 43% (Fig. 6E). Consistent with the first GCAUG’s stronger regulatory effect, RNA fragments encompassing the first GCAUG are isolated in higher yield than the second, possibly indicating a higher affinity for the Rbfox1/LASR complex. Deleting both motifs together brought the PSI down to 3%, indicating that although the first GCAUG exerts a stronger regulatory effect, they contribute additively to exon regulation. We also deleted the first G-rich region along with both GCAUG motifs. This led to complete exon skipping, mirroring the splicing observed in Rbfox deficient and Rbfox(F125A) expressing cells. Thus, hnRNP H/F and Rbfox1, together, provide the splicing enhancement needed to include this *CAMKK2* exon.

We also assessed the regulatory effects of motifs in protected sites adjacent to an activated exon in *MARK3* (Fig. S10). This exon was activated in both the wildtype and F125A expressing cells, with a slightly higher PSI in the F125A (80%) compared to wildtype (66%). The downstream intron of this exon contains three protected sites: the most proximal region 1 contains G-rich and CU-rich elements, the middle region 2 contains GCAUG and CU-rich elements, and the distal region 3 contains GCAUG and GU-rich elements (Fig. S10A). The gene segment containing this exon, along with 500 nucleotides each of upstream and downstream intronic sequences, was cloned into the same DUP mini-gene backbone used for the *CAMKK2* exon 16 (Fig. S10B). The splicing pattern of the *MARK3* mini-gene largely recapitulates the splicing of the endogenous gene. The exon is included in both the wildtype and F125A expressing cells and the PSI is higher in the F125A expressing cells, 31%, compared to the wildtype, 16% (Fig. S10C).

We then deleted potential regulatory motifs in protected sites and tested their effects on splicing in cells expressing wildtype Rbfox1 (Fig. S10D-E). Some of the elements such as the G-rich element in region 1 and the CU-rich element in region 2 had almost no effect on the inclusion of this exon, yielding an PSI of 17% and 14% respectively. In contrast, deleting the GCAUG elements in either region 2 or 3 reduced inclusion of this exon (PSI of 8% and 9% respectively), demonstrating that interaction of Rbfox1/LASR with these elements contributes to the regulation of this exon. Interestingly, mutation of the CU-rich element in region 1 had the strongest effect on splicing, yielding an PSI of 2%. The splicing of these mutants was also assessed in cells expressing Rbfox1(F125A) (Fig. S10F). Deletion of the CU-rich motif in region also had the strongest effect in these cells, resulting in a PSI of 1%. Interestingly, the mini-genes lacking the GU-rich element in region 1 or the GCAUG in region 2 both exhibited slightly higher PSIs, 44% and 49% respectively. Thus, for this exon that is regulated by both the wildtype and the F125A Rbfox1, the element with the strongest stimulatory effect is not a Rbfox binding site but rather the CU-rich element.

## Discussion

### Recognition of multi-element RNA modules by the Rbfox/LASR complex

We previously found that most of the Rbfox protein in the nucleus is associated with nascent RNA and bound to LASR, a protein complex containing hnRNP M, hnRNP H/F, hnRNP C, Matrin3, hnRNPUL2, NF110/NFAR-2, NF45, and DDX5 (Damianov et al. 2016; Ying et al. 2017). We further found that Rbfox binding to GCAUG elements and its splicing activity were affected by the LASR subunit hnRNP M (Damianov et al. 2016). However, the RNA contacts made by the Rbfox/LASR, beyond those mediated by Rbfox’s RRM, were not clear. We show here that isolated Rbfox1/LASR complexes contain small fragments of nuclease-protected RNA. These RNAs contain the Rbfox binding motif GCAUG, as well as binding motifs for the LASR subunits hnRNP M, hnRNP H/F, hnRNP C, and Matrin3. RNA fragments containing a UAG motif were also isolated. A well-known binder to this motif hnRNP A1 is not present in LASR. This motif may interact with LASR subunits whose RNA binding properties are not completely characterized, including NF45, NF110, and DDX5. Alternatively, one of the LASR subunits may have a secondary binding specificity in addition to its known interacting motifs. Future *in vitro* binding assays and/or CLIP analysis can elucidate the additional binding preferences of the LASR subunits.

Our IP-seq *in vivo* footprinting approach for identifying RNA-protein interactions in cells, yields different information than CLIP protocols. CLIP is applied under stringent immunoprecipitation conditions aimed at isolating a single protein after crosslinking in cells (Hafner et al. 2021). Sequencing the isolated fragments after reverse transcription identifies the sites of crosslinking on the RNA by either mutations introduced during reverse transcription or by the position of a reverse transcriptase stop. These sites of crosslinking are not footprints and do not provide information on the amount of RNA engaged by the isolated protein, although specific motifs recognized by the protein can often be identified by their enrichment at the crosslinking sites. Many direct RNA interactions can be identified by CLIP, but individual proteins and RNA-protein interactions can vary widely in the efficiency of their crosslinking and recovery in the immunoprecipitate. Also, some proteins may crosslink to nucleotides that are not engaged in a specific protein interaction but are simply nearby in space. In the eCLIP data for HEK293 cells, Rbfox2 crosslinks not just to GCAUG but across the regions surrounding these elements. This pattern may result from crosslinking due to proximity, where Rbfox is close to these elements because it is bound to LASR subunits that interact with these regions. Alternatively, other proteins in LASR may be crosslinking and remain present in the immunoprecipitated material to yield crosslinked RNA reads. hnRNPH/F migrate close to Rbfox in SDS-PAGE. If these proteins co-immunoprecipitate with Rbfox under CLIP conditions, the RNAs crosslinked to them will likely be excised from the gel along with the RNA crosslinked to Rbfox and sequenced. A recent report suggests that protein co-factors can contribute to the background of RNA fragments detected in some CLIP studies (Guo et al., 2024).

Our IP-seq method requires that the interaction of a protein or larger complex with the RNA be sufficiently stable. It is expected that RNAs engaged in interactions that undergo exchange in solution will not survive the nuclease treatment. Such interactions might be captured by CLIP. So far, we have used IP-seq to identify transcriptomic interactions of the U2 snRNP and the Rbfox1/LASR complex. It remains to be seen how many other protein/RNA interactions can be studied in this way. IP-seq allows analysis of RNA interactions with intact protein complexes. This means that one can’t always determine which protein in the complex is making a direct contact, without testing mutant proteins that eliminate particular interactions. This strategy allowed us to distinguish the RNA contacts of Rbfox1 from those of LASR subunits. It will be interesting to extend this analysis to mutations in LASR proteins. However, Rbfox is unusual in that a single point mutation is known to dramatically reduce RNA binding. The other proteins in LASR have multiple RRM or other domains and identifying suitable mutations that prevent RNA binding while retaining LASR assembly will require extensive analysis.

Complexes of transcription factors often bind to elements of DNA arranged in a particular order (Whitington et al. 2011). The structures of these complexes constrain how their DNA binding proteins contact their target sequence. To examine whether there were optimal arrangements of motifs in RNA that might facilitate recruitment of Rbfox/LASR, we analyzed the positional distributions of the different Rbfox/LASR elements relative to the GCAUG motifs. We find that motifs for LASR subunits are enriched within 1-10 nucleotides of the Rbfox motif. GU-rich motifs are often directly abutting the GCAUG, while G-rich elements are usually slightly further away. The GU-elements are more commonly upstream of the Rbfox site and G-rich elements more commonly downstream. However, these were not strong preferences and GU elements and G-rich elements can both be found on either side of the GCAUG. Although these preferences hint at a spatial arrangement of hnRNP H/F and hnRNP M relative to Rbfox, the arrangements are more variable than sometimes seen for transcription factors. The greater flexibility of single stranded RNA compared to double stranded DNA might accommodate a wider range motif arrangements. Moreover, hnRNP M and hnRNP H/F each have multiple RNA binding domains that could be positioned to allow binding of motifs at different locations relative to the Rbfox motif.

Splicing of the mutually exclusive exons 5A and 5B in the FGFR gene of *C. elegans* egl-15 is controlled by a UGCAUGGUGUGC element. The GCAUG in this sequence is bound by the by the *C. elegans* Rbfox paralogue ASD-1 and the GUGUGC element is recognized by the protein SUP-12 (Kuwasako et al., 2014). The structure of a ternary complex containing the RNA binding domains of ASD-1 and SUP-12 bound with the UGCAUGGUGUGC sequence shows that the two proteins simultaneously contact this RNA and sandwich the G7 nucleotide between their binding domains. SUP-12 does not have direct paralogues in mammals. However, hnRNP M is a well-known binder to motifs with repeating GU nucleotides. The presence of GCAUG elements directly adjacent to GU-rich elements in both *C. elegans* and humans suggests an evolutionary conserved splicing regulatory module. Understanding the similarity of SUP-12 and hnRNP M interactions will require future structural studies.

We found that LASR in complex with the mutant Rbfox1(F125A) lost binding to almost all binding sites that contained GCAUG. Although it is expected that the F125A mutation would reduce Rbfox’s affinity for these sites, resulting in loss of protection, many of these sites also contained motifs for LASR. The affinity of LASR subunits for these sites may not be sufficient for protection from the nuclease cleavage. Rbfox likely drives LASR to bind to motifs in these regions through its highly specific and strong interaction with GCAUG. LASR, in turn, specifies Rbfox’s binding to GCAUGs that are adjacent to its own motifs. Rbfox/LASR target recognition seems to balance the highly specific interaction of Rbfox with more versatile recognition of RNA by LASR. Furthermore, many Rbfox1/LASR protected sites lack a GCAUG element but contain tandem GU-rich or G-rich elements. These tandem elements presumably increase the RNA’s affinity for the multiple RNA binding domains present in hnRNP M and hnRNP H/F.

### Regulation of alternative pre-mRNA splicing by combinatorial interactions between Rbfox and LASR co-factors

Exons carrying Rbfox binding motifs are part of large coordinated splicing programs in neurons, muscle, and other tissues (Conboy 2017). Such splicing programs include splicing factors that cross regulate other RNA-binding proteins, creating complex regulatory networks (Jangi et al. 2014; Ule and Blencowe 2019). The Rbfox/LASR complex presents an interesting example of how multiple factors might combine to regulate particular exons. Previously, we found that the LASR subunit hnRNP M can affect a subset of Rbfox targets (Damianov et al. 2016). But it was not clear how other LASR subunits might affect Rbfox’s regulatory activity. Here, we find that exons activated by Rbfox1 carry adjacent binding motifs for multiple LASR subunits. These LASR motifs can co-occur with Rbfox motifs within a binding site, thereby influencing Rbfox binding to a regulatory region. Mini-gene experiments show that motifs for LASR subunits can directly influence activation of exons that are also dependent on Rbfox motifs.

In addition to binding to exact GCAUG motifs, we find that Rbfox also activates splicing through secondary motifs adjacent to LASR elements. A previous study found that increased Rbfox levels during neuronal development may enhance its binding to these secondary motifs (Begg et al. 2020). We show that beyond increased site occupancy due to higher concentrations, association with LASR co-factors can also influence Rbfox binding to secondary motifs. This association can thus broaden Rbfox targeting to an additional set of exons. It will be interesting to examine whether particular physiological processes are controlled via the combined Rbfox secondary motifs and LASR motifs.

All three mammalian *RBFOX* genes produce an alternatively spliced protein isoform that lacks the second half of the RNA Recognition Motif (RRM). The expression of this Rbfox(ΔRRM) is auto-regulated by full-length Rbfox, which binds to a GCAUG element upstream of an RRM-encoding exon to cause exon skipping (Damianov & Black, 2010). This isoform produces a stable protein that appears to act in a dominant negative manner, counteracting regulation by the full-length isoform. Here, we find that when Rbfox’s affinity for GCAUG is reduced by a point mutation, it alters binding of the LASR complex across the transcriptome. LASR in complex with the F125A Rbfox1 mutant loses binding to some sites, retains binding to others, and shows enhanced binding to a more limited set of sites. It will be interesting to perform the IP-seq of LASR bound with Rbfox(ΔRRM) to assess whether this complex also binds to novel sites. Through its interaction with LASR, the ΔRRM isoform might exhibit activities beyond acting as a dominant negative factor.

Many mutagenesis analyses of splicing regulatory elements have found that multiple elements adjacent to a cassette exon can exert additive effects on exon inclusion or skipping (Han et al. 2005; Modafferi and Black 1999; Ryan and Cooper 1996; Smith and Valcárcel 2000). It is usually not clear from these studies whether these elements are being bound by multiple independent factors or even if these factors are simultaneously binding the same RNA. Our data identify closely spaced multipart elements on the same RNA fragment bound by Rbfox1/LASR. Multipart elements can also be more widely spaced within a regulatory region and give rise to separate protected RNAs. It is not known how many unit complexes of Rbfox/LASR interact with these dispersed regulatory sequences. The Rbfox/LASR complex can multimerize into higher-order structures through the Rbfox C-terminal domain, and this multimerization is required for Rbfox dependent activation of many exons (Ying et al. 2017). Thus, each discrete Rbfox/LASR particle bound at protected site in a regulatory region may be part of a larger multimerized complex. Such multimerized complexes could bridge large segments of intronic RNA and allow more distal elements to affect splicing of an exon. In the future, it will be interesting to assess how the loss of multimerization through mutation affects the pattern of binding by Rbfox/LASR and its activity in splicing.

## Materials and Methods

### Cell culture conditions

Cells were cultured in 90% DMEM ([+] 4.5 g/L glucose, L-glutamine[-] sodium pyruvate, Corning) and 10% (v/v) fetal bovine serum (Omega scientific) at 37 °C with 5% CO2. Mycoplasma contamination was monitored using the PCR-based VenorGeM® Mycoplasma Detection Kit.

### Cell lines

Flp-In™ T-REx™ 293 Cell Line (ThermoFisher Scientific) is the parental line for all derived cell lines. As previously described, a Rbfox-deficient line was derived from the parental line by CRISPR/Cas9 deletion of the first constitutive Rbfox2 exon (Damianov et al. 2016; Ying et al. 2017). FLAG-Rbfox1(WT) and FLAG-Rbfox1(F125A) were integrated into the FRT site and a mixed population of each respective line was selected via hygromycin treatment. Cells were treated for 48 hours with doxycycline to induce expression of the transgenes.

### Plasmid construction

All the primers can be found in Table 1.

**Table 1.**
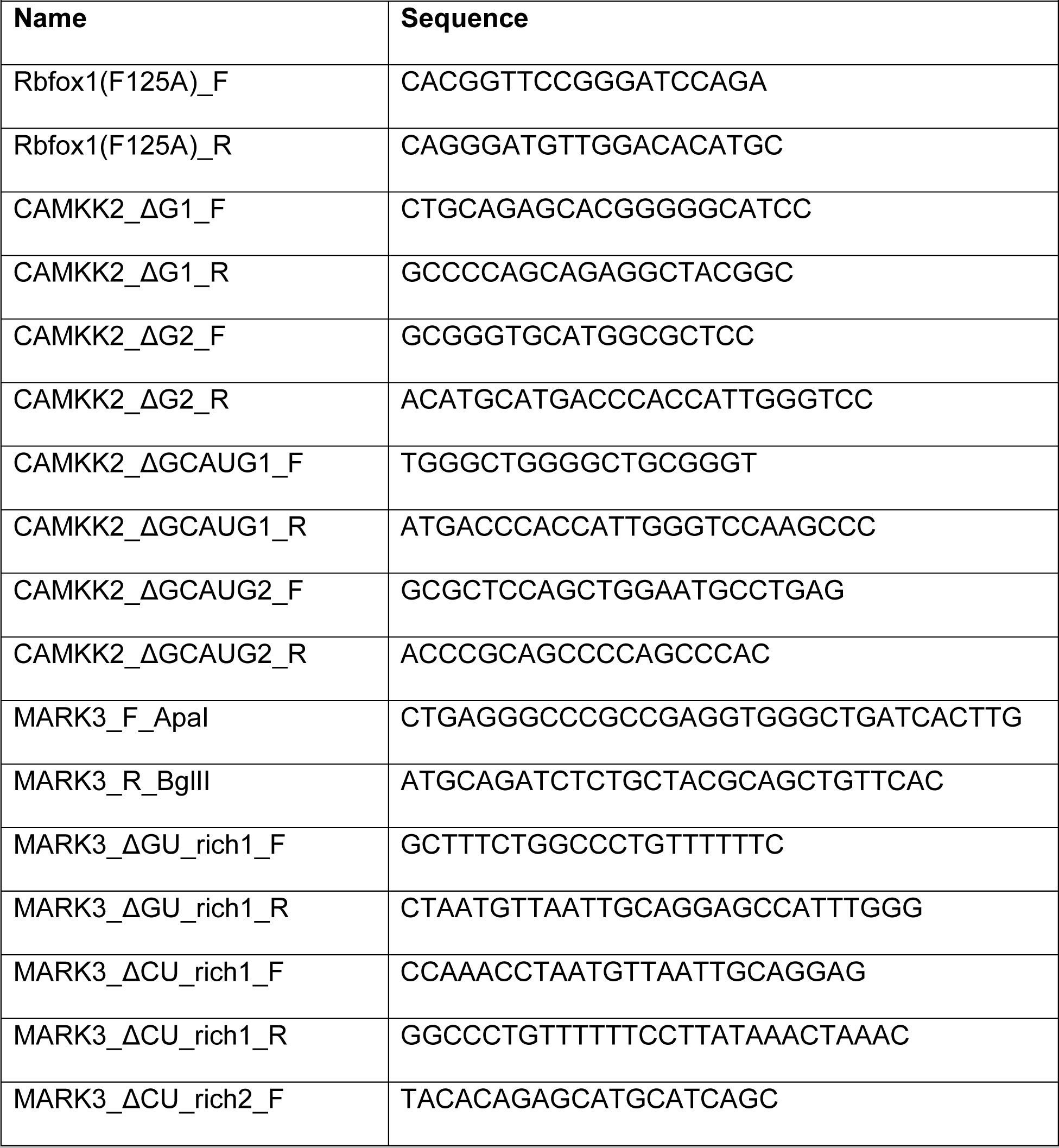
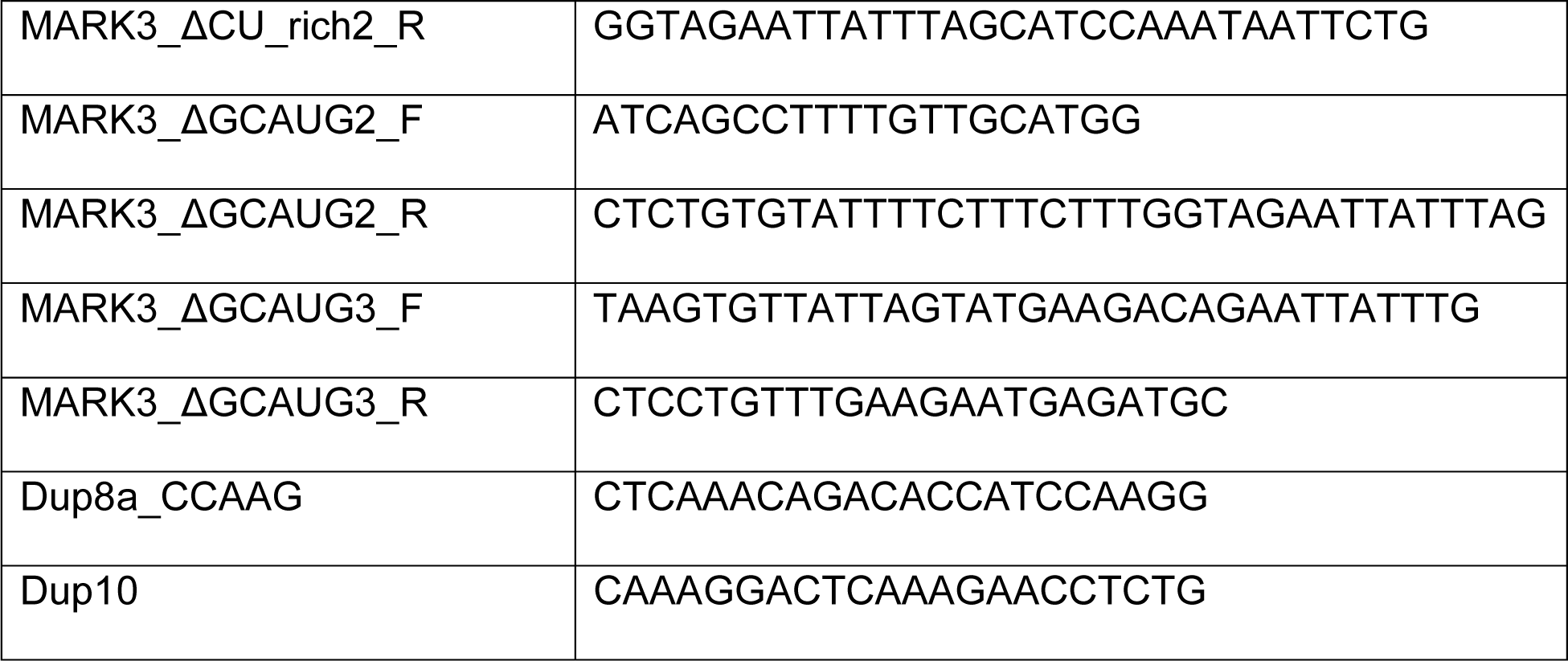
Oligos.

FLAG-Rbfox1(WT): Rbfox1 with an N-terminal 1x FLAG tag was cut out of pcDNA3.1 and ligated into the pcDNA™5/FRT/TO vector via restriction cloning using HindIII-HF and XhoI sites.

FLAG-Rbfox1(F125A): Based on the study by Auweter et al., a critical residue in Rbfox’s RRM is phenylalanine 126 in the Swissprot Q9NWB1 entry. This phenylalanine — underline here, VSNIP<u>F</u>RFRD **—** is at position 125 in our Rbfox1 construct which lacks a single amino acid in the N-terminus when aligned with the Q9NWB1 construct. We mutated this phenylalanine to an alanine in the pcDNA5/FRT/TO FLAG-Rbfox1(WT) vector via site-directed mutagenesis PCR using primers Rbfox1(F125A)_F and Rbfox1(F125A)_R.

CAMKK2 exon 16 splicing reporter: the CAMKK2 exon 16 splicing reporter construct used by Cao et al. was obtained from Jiuyong Xie. The region of this construct containing *CAMKK2* was subcloned into the DUP-51M1 backbone by restriction cloning via BglII and ApaI sites. This backbone has all GCAUG motifs and potential hnRNP M sites mutated (Damianov et al. 2016). This DUP-51M1 CAMKK2 exon 16 construct was used as the splicing reporter in this study. Deletions of regulatory elements in figure 4 were done on this mini-gene by site-directed mutagenesis via the following primers: ΔG1 = CAMKK2_ΔG1_F + CAMKK2_ΔG1_R, ΔG2 = CAMKK2_ΔG2_F + CAMKK2_ΔG2_R, ΔGCAUG1 = CAMKK2_ΔGCAUG1_F + CAMKK2_ΔGCAUG1_R, ΔGCAUG2 = CAMKK2_ΔGCAUG2_F + CAMKK2_ΔGCAUG2_R.

Mark3 exon 16 splicing reporter: a region comprising exon 16 along with 522 nucleotides in the upstream and 539 nucleotides in the downstream intron was PCR amplified with the following primers containing ApaI and BglII cut sites: MARK3_F_ApaI, MARK3_R_BglII. This amplified region was cloned into the DUP-51M1 backbone via restriction cloning using the BglII and ApaI sites. Regulatory elements shown in Supplementary Figure 10 were deleted via site-directed mutagenesis by the following PCR primers: ΔGU-rich1 = MARK3_ΔGU_rich1_F + MARK3_ΔGU_rich1_R, ΔCU-rich1 = MARK3_ΔCU_rich1_F + MARK3_ΔCU_rich1_R, ΔCU-rich2 = MARK3_ΔCU_rich2_F + MARK3_ΔCU_rich2_R ΔGCAUG2 = MARK3_ΔGCAUG2_F, MARK3_ΔGCAUG2_R, ΔGCAUG3 = MARK3_ΔGCAUG3_F, MARK3_ΔGCAUG3_R.

### Purification of ribonucleoprotein complexes

Rbfox/LASR complexes were purified as previously described (Damianov et al. 2016). To isolate nuclei, cells were grown to 80-90% confluency and harvested. Cell pellets were resuspended in nine volumes of ice-cold homogenization buffer (10 mM HEPES-KOH pH 7.6, 15 mM KCl, 1 mM EDTA, 1.8 M sucrose, 5% Glycerol, 0.15 mM Spermine, 0.5 mM Spermidine) and homogenized with the gentleMACS homogenizer. The homogenate was overlaid onto 10 mL of ice-cold cushion buffer (10 mM HEPES-KOH pH 7.6, 15 mM KCl, 1 mM EDTA, 2.0 M sucrose, 10% Glycerol, 0.15 mM Spermine, 0.5 mM Spermidine) in a SW32Ti ultracentrifugation tube and centrifuged at 28,100 rpm (96,970 g) for 1 hr at 4°C.

To lyse nuclei and obtain an extract from the chromatin containing pellet, supernatant and cushion buffer were discarded and the pelleted nuclei were suspended in 1 mL of nuclear resuspension buffer (10 mM HEPES-KOH pH 7.6, 15 mM KCl, 1 mM EDTA, 0.15 mM spermine, 0.5 mM spermidine) and aliquoted into three separate tubes of equal volume. The samples were centrifuged for 5 mins at 1k rcf, the supernatant was discarded, and the nuclei were resuspended in at least 10x volume of nuclear lysis buffer (20 mM HEPES-KOH pH 7.6, 150 mM NaCl, 1.5 mM MgCl2, 0.5 mM DTT, 1x protease inhibitors, and 0.6% Triton X-100) and kept on ice for 5 mins for nuclear lysis.

This lysate was then centrifuged at 20k rcf for 10 mins and the supernatant was kept as the nucleoplasm portion. The same volume as the nucleoplasm portion of the nuclear lysis buffer was added to the pellet and Benzonase nuclease was added to all nucleoplasm and pellet-containing samples to a final concentration of 5 units/ul. The nuclease digestion was done until the pellet could be resuspended by a P200 ul tip. The nuclease treated samples were then centrifuged at 20k rcf for 10 mins and the supernatant was kept and the pellet was discarded.

To perform immunoprecipitations, the supernatant was added to 7.5 ul of packed M2 FLAG agarose beads (Sigma) and this mixture was rotated overnight at 4°C. The beads were washed 5x with each wash containing 1 mL of wash buffer (20 mM HEPES-KOH pH 7.6, 150 mM NaCl, and 0.05% Triton X-100). 50 ul of elution buffer (20 mM HEPES-KOH pH 7.9, 150 mM NaCl, and 150 μg/ml of 3xFLAG peptide) was then added to the beads and this mixture was agitated intermittently (15 sec on, 4:45 min off) at 1100 rpm at 4°C for 1 hr. Supernatant was saved as eluate for further processing.

### SDS-PAGE analysis of purified complexes

For SDS-PAGE analysis of immunoprecipitated Rbfox1/LASR complexes, 20% of the immunoprecipitated material was denatured and run on a NuPAGE® Novex 4-12% Bis-Tris gel. The gel was then stained with SYPRO Ruby (ThermoFisher Scientific) overnight following the manufacturer’s instructions. Stained gels were imaged with the Amersham Typhoon.

### Urea-PAGE analysis and sequencing of nuclease-protected RNA

For all phenol-chloroform extractions, the aqueous phase was separated from the organic phase using Phase Lock Gel Heavy tubes (QuantaBio). All ethanol precipitations were done overnight at −20°C and Glycoblue was used. 40% of the immunoprecipitated material was deproteinized with Proteinase K and an acid-phenol chloroform extraction followed by ethanol precipitation was done. This material was then DNAse treated and dephosphorylated in a one-pot reaction containing TurboDNAse and FastAP Thermosensitive Alkaline Phosphatase. Another Proteinase K digestion followed by phenol-chloroform extraction and ethanol precipitation was then done on this material. 5% of the resulting material was used for end-labeling and the rest was used to make sequencing libraries. T4 PNK was used to ligate γ-[32P] ATP (Perkin Elmer) to 5’ end of the RNA. This material was denatured with formamide and run on a 10% Urea-PAGE. The gel was dried and used to generate an autoradiograph which was imaged with the Amersham Typhoon. The other 95% of the material was used to prepare sequencing libraries using a modified iCLIP library preparation protocol (Damianov et al. 2024). This protocol was followed as stated except for the U2 snRNA degradation. This library was sequenced with the NovaSeq 6000 SP 2×100bp.

### Sequencing RNA for splicing analysis

Total RNA was extracted using Trizol. This RNA was purified via Zymo’s RNA Clean & Concentrator-5 Kit with a DNAseI treatment step. cDNA was prepared from poly-A selected RNA using the Illumina TruSeq kit. This cDNA was sequenced in one lane of the NovaSeq X.

For assessing expression of Rbfox proteins in the RNA-seq experiment, 50% of harvested cells were lysed with RIPA. Total protein concentrations were measured with Pierce BCA kit and the same amount of total protein was denatured and analyzed per sample with 10% SDS-PAGE. Immunoblotting was done by transferring the samples to a PVDF membrane and probing with FLAG and GAPDH primary antibodies and fluorescent secondary antibodies. All antibodies used can be found in Table 2.

**Table 2.** Antibodies.

|  |  |  |
| --- | --- | --- |
| FLAG | MilliporeSigma | F3165 |
| GAPDH | Invitrogen | PA1-987 |
| hnRNPH/F | Santa Cruz Biotechnology | sc-32310 |
| Anti-mouse Cy5 | Cytiva | PA45009 |
| Anti-rabbit Cy3 | Cytiva | 28-9011-06 |

### Mini-gene analysis

All transfections and the subsequent steps were done in triplicate for all the constructs tested. A mixture of lipofectamine 2000 and the construct of interest was added to cells in complete growth medium and incubated for 6 hrs at 37 °C. The media was then changed to fresh complete growth medium and cells were incubated and harvested the next day after 24 hrs of being transfected. Total RNA was extracted as described in the “RNA-sequencing for splicing analysis” section. 4.5 ug of this RNA was reverse transcribed via SuperScript™ (Invitrogen) and an oligo with 20 dT nucleotides. cDNA was PCR amplified for 15 cycles using Dup8a_CCAAG and Dup10 primers via the GoTaq Green Master Mix (Promega). Amplified material was run on a 2.5% agarose gel, stained with SYBR Safe, and imaged via the iBright Imaging System (ThermoFisher Scientific). ImageJ was used to calculate the intensity of the bands. Percent spliced in (PSI) values were calculated for reach replicate by dividing the intensity of the upper band by the combined intensity of both bands, and the average was calculated.

### siRNA knockdown of hnRNPH/F

For siRNA knockdowns, siHNRNPH/F or siControl (Silencer Select Negative Control #1 from Thermo Fisher) were transfected into cells with Lipofectamine RNAiMAX (Invitrogen) per manufacturer’s protocol. The sequence for siHNRNPH/F is GGAAGAAAUUGUUCAGUUC (Garneau et al. 2005). After 5 hours, medium was changed to complete growth medium and cells were incubated for 24 hours. Another transfection was then done with siHNRNPH/F or siCONTROL using lipofectamine 2000 with both conditions including the DUP-51M1 CAMKK2 exon 16 construct. After 5 hours, medium was changed to complete growth medium and cells were incubated for 48 hours. Cells were subsequently harvested and split into two equal portions to be used for RNA extraction and RIPA lysis followed by immunoblotting.

### Mapping IP-seq sequencing data and defining clusters

Samples were demultiplexed, PCR duplicates were removed, and the reads were mapped to hg38 using STAR (Dobin et al., 2013). YODEL was used to define clusters in regions containing at least 10 reads in merged replicates (Palmer et al. 2017). RPKM values of clusters were calculated with the SeqMonk software (v1.45.4, Babraham Institute). A chi-squared test was performed comparing RPKM of size-matched clusters from the experimental to the control IP-seq clusters. All clusters with an FDR < 0.05 and log_2_(sample/control) > 0 were called as significant and further processed.

### Annotation of clusters

Clusters were annotated based on Ensembl Canonical genes annotations. Based on the annotations, the percent of clusters that fell into different gene types and regions were determined. To define clusters in introns of protein-coding genes, intronic sites that also resided in regions annotated as snRNA, miRNA, scaRNA, snoRNA, ncRNA, and lncRNA in Ensembl canonical genes or NCBI Refseq databases were discarded.

### Motif enrichment analysis

Reads that fall into clusters in introns of protein-coding introns were extracted. A background set was generated by sampling random regions from introns that contained the clusters. HOMER was used for motif analysis via the following command: findMotifsGenome.pl <experimental set> hg38 <output file> -bg <background set> -rna - len 4,5,6 -S 10 -size given (Heinz et al. 2010).

Enrichment of pentamers and hexamers was determined by comparing their frequency in clusters within introns of protein-coding genes to randomly sampled regions from introns that contain clusters. Z-scores were determined for these pentamers and hexamers as previously described (Damianov et al. 2016).

### Determining protected vs unprotected GCAUGs

The SeqKit software was used to calculate coordinates for all (U)GCAUG, UGCAUG, and GCAUG (without a 5’ U) elements (Shen et al., 2016). Expressed elements were defined as those present in introns that contained at least one IP-seq cluster. We then determined the protection status of these elements by assessing their overlap with IP-seq clusters. The percent of protected versus unprotected elements was calculated accordingly.

### Motif co-occurrence analysis

For quantifying how many protected fragments containing GCAUGs also contained another motif, protected fragments from clusters in introns of protein-coding genes were annotated with HOMER. The co-occurrence of GCAUG with other motif types was then determined based on this annotation. Furthermore, these annotated protected fragments were used to generate plots of the distribution of length of reads associated with different types of motifs.

To determine co-occurrence of GCAUGs with other motifs, a region of 50 nucleotides upstream and 50 nucleotides downstream of each GCAUG was defined. To generate plots displaying the positional frequency of motifs around GCAUGs, GCAUG was considered to be at position 0. If a motif was found starting at 1 nucleotide downstream of a GCAUG it was considered to be at position 1. Conversely, if a motif was found to end at 1 nucleotide upstream of a GCAUG, it was considered to be at position −1. The number of times a motif occurred at each position was then determined relative to GCAUGs for all motifs in each motif category. Then the total number of times a motif occurred at a given position was divided by the total number of GCAUGs to determine the fraction of regions with a given motif type. A plot was then generated to display this fraction for different motif types surrounding unprotected and protected GCAUGs separately.

### Differential binding analysis via DESeq2

All significant clusters of Rbfox1(WT)/LASR and Rbfox1(F125A)/LASR were merged. To avoid merging overlapping clusters into a single bigger cluster, the clusters were shortened to only include the region that is covered by 50% of the total reads within clusters. DEseq2 analysis was performed on this merged set of clusters comparing wildtype to F125A IP-seq (Love et al. 2014). Differentially bound sites were defined as the following: WT preferential sites have a log2FoldChange (WT/FA) >=1 and FDR < 0.05, FA preferential sites have a log2FoldChange (WT/FA) <= −1, common sites have a log2FoldChange (WT/FA) > −0.1 and < 0.1.

### Splicing analysis of sequenced RNA via rMATS

Alternative splicing was analyzed by rMATS Turbo (Wang et al. 2024). Pairwise comparisons were done for WT vs KO, FA vs KO, and WT vs FA. Events that had at least 5 junction reads in each replicate were filtered for further analysis. WT-regulated exons are those that have a |ΔPSI(WT-KO)| >= 0.15, FDR <= 0.05, and |ΔPSI(FA-KO)| <= 0.05. WT/FA-regulated exons have a |ΔPSI(WT-KO)| >= 0.15, |ΔPSI(FA-KO)| >= 0.15, | ΔPSI(WT-KO) - ΔPSI(FA-KO)| <= 0.15, and a FDR < 0.05 in both comparisons. FA-regulated exons have |ΔPSI(FA-KO)| >= 0.15, FDR <= 0.05, and |ΔPSI(WT-KO)| <= 0.05.

### Mapping clusters around regulated exons

Upstream and downstream intronic regions were defined by the region spanning the cassette exon and its adjacent exons as determined by rMATS. Clusters that occurred in these regions were extracted and split according to the elements they contained. The occurrence of clusters containing elements of interest were assessed at every single nucleotide in both the upstream and downstream introns. The total number of clusters that occurred at a given nucleotide were summed and divided by the total number of exons to give the fraction of regions containing clusters with an element of interest. A bootstrapping strategy described by Yee et al. was used to assess the enrichment of clusters around regulated exons compared to their frequency around unregulated exons (Yee et al. 2019).

## Data availability

The IP-seq, YODEL defined clusters, RNA-seq, and rMATS tables are available in the GSE272026 entry at GEO.

## Acknowledgements

We thank Jiuyong Xie for providing the CaMKK2 exon 16 splicing reporter minigene and Andrey Damianov for the DUP-51M1 backbone. We thank Andrey Damianov and all the members of the Black Laboratory for their help and constructive discussions. This work was supported by NIH grants R35GM136426 and R21HG012624 to D.L.B., and a SEED grant from the Jonsson Comprehensive Cancer Center at UCLA to D.L.B. P.P. was supported by T32GM008042, T32GM152342, and NIMH F30MH130075.

## Author Contributions

P.P. and D.L.B. conceived and designed the research; P.P. performed experiments with help from K.O.; P.P. and C.-H.L. performed bioinformatics analysis; P.P. and D.L.B. wrote the manuscript with input from other co-authors.

